# Influenza A virus NS1 effector domain is required for PA-X mediated host shutoff

**DOI:** 10.1101/2023.10.02.560421

**Authors:** Juliette Bougon, Eileigh Kadijk, Lucie Gallot-Lavallee, Bruce Curtis, Matthew Landers, John M. Archibald, Denys A. Khaperskyy

**Author notes:** These authors contributed equally (Juliette Bougon, Eileigh Kadijk).

## Abstract

Many viruses inhibit general host gene expression to limit innate immune responses and gain preferential access to the cellular translational apparatus for their own protein synthesis. This process is known as host shutoff. Influenza A viruses (IAVs) encode two host shutoff proteins: nonstructural protein 1 (NS1) and polymerase acidic X (PA-X). NS1 inhibits host nuclear pre-messenger RNA maturation and export, and PA-X is an endoribonuclease that preferentially cleaves host spliced nuclear and cytoplasmic messenger RNAs. Emerging evidence suggests that in circulating human IAVs NS1 and PA-X co-evolve to ensure optimal magnitude of general host shutoff without compromising viral replication that relies on host cell metabolism. However, the functional interplay between PA-X and NS1 remains unexplored. In this study, we sought to determine if NS1 function has a direct effect on PA-X activity by analyzing host shutoff in A549 cells infected with wild type or mutant IAVs with NS1 effector domain deletion. This was done using conventional quantitative reverse transcription polymerase chain reaction techniques and direct RNA sequencing using nanopore technology. Our previous research on the molecular mechanisms of PA-X function identified two prominent features of IAV infected cells: nuclear accumulation of cytoplasmic poly(A) binding protein (PABPC1) and increase in nuclear poly(A) RNA abundance relative to the cytoplasm. Here we demonstrate that NS1 effector domain function augments PA-X host shutoff and is necessary for nuclear PABPC1 accumulation. By contrast, nuclear poly(A) RNA accumulation is not dependent on either NS1 or PA-X mediated host shutoff and is accompanied by nuclear retention of viral transcripts. Our study demonstrates for the first time that NS1 and PA-X effects on host gene expression are not simply additive and that these factors functionally interact in mediating host shutoff.

**IMPORTANCE:** Respiratory viruses including influenza A virus continue to cause annual epidemics with high morbidity and mortality due to limited effectiveness of vaccines and antiviral drugs. Among the strategies evolved by viruses to evade immune responses is host shutoff – a general blockade of host messenger RNA and protein synthesis. Disabling influenza A virus host shutoff is being explored in live attenuated vaccine development as an attractive strategy for increasing their effectiveness by boosting antiviral responses. Influenza A virus encodes two proteins that function in host shutoff: the non-structural protein 1 (NS1) and the polymerase acidic X (PA-X). We and others have characterised some of the NS1 and PA-X mechanisms of action and the additive effects that these viral proteins may have in ensuring the blockade of host gene expression. In this work we examined whether NS1 and PA-X functionally interact and discovered that NS1 is required for PA-X to function effectively. This work significantly advances our understanding of influenza A virus host shutoff and identifies new potential targets for therapeutic interventions against influenza and further informs development of improved live attenuated vaccines.

## INTRODUCTION

Influenza A viruses (IAVs) are enveloped viruses with a negative sense RNA genome divided into 8 segments [1]. IAVs are important human and animal pathogens with high pandemic potential. The introduction of new IAV strains from zoonotic reservoirs into the human population have led to a number of pandemics throughout history and this threat continues today [2,3]. Between pandemics, IAVs that circulate in humans continue to be responsible for annual epidemics worldwide that cause significant morbidity and mortality. Previous infections or vaccinations do not offer complete protection because the virus evades adaptive immunity by constantly changing its major epitopes for neutralizing antibodies – a process called antigenic drift [3]. In the absence of virus-neutralizing adaptive immunity, innate immune responses represent an important first line of defence against viruses. Eukaryotic cells are capable of recognising infection by detecting pathogen associated molecular patterns (PAMPs) through an array of sensors [4]. The most important sensor for IAV and other negative sense RNA viruses in infected cells is the retinoic acid inducible gene I (RIG-I) [5,6]. RIG-I is an RNA helicase that recognises viral genomic RNA with 5’ triphosphate ends. Upon viral RNA binding, RIG-I changes conformation, undergoes a series of posttranslational modifications and oligomerizes together with the mitochondrial antiviral signaling protein (MAVS) to initiate an activation cascade that culminates in transcriptional induction of type I interferon (IFN) and other antiviral cytokines [5]. These cytokines signal through their receptors to induce an antiviral state in infected and neighbouring cells and modulate responses by the immune system. Specifically, type I IFN exerts a potent antiviral effect through induction of an array of IFN-stimulated genes (ISGs) that interfere with various aspects of viral replication [4].

To counteract innate antiviral responses, IAV evolved multiple strategies to interfere with the sensing of viral nucleic acids and downstream signalling from RIG-I/MAVS [7–9]. Furthermore, to ensure efficient suppression of antiviral responses, IAV executes host shutoff – a general inhibition of host gene expression in infected cells. In addition to blocking expression of IFNs and ISGs, host shutoff facilitates access to cellular translation machinery by viral messenger RNAs (mRNAs) by alleviating competition from cellular transcripts [10]. Many RNA and DNA viruses encode host shutoff factors. One important type of these factors is the nucleases that function through cleavage and degradation of host mRNAs [11]. These include the virion host shutoff (VHS) protein of herpes simplex virus-1 (HSV-1) [12,13], the SOX endonuclease of Kaposi’s sarcoma-associated herpes virus (KSHV) [14,15], and the non-structural protein 1 (NSP1) of severe acute respiratory syndrome coronaviruses (SARS-CoV and SARS-CoV2) [16,17]. IAV also encodes a host shutoff endonuclease polymerase acidic X protein (PA-X) [18,19]. PA-X is a highly conserved protein produced via +1 ribosomal frameshifting at Phe-191 during translation of the segment 3-derived mRNA, which encodes the PA subunit of the viral RNA-dependent RNA polymerase (RdRp) [18,20,21]. Although relatively modest amounts of PA-X accumulate in IAV-infected cells due to low frameshifting efficiency, PA-X is a very potent host shutoff protein and the main RNA degradation factor [18,22]. Recombinant viruses with an altered frameshifting site in the PA gene that prevents PA-X production are less effective at blocking expression of antiviral and proinflammatory cytokines in cell culture models and in *in vivo* mouse models of infection [18,23–25]. Our previous work examining molecular mechanisms of PA-X host shutoff revealed that this endonuclease selectively targets host RNA polymerase II transcripts and that its activity results in depletion of cytoplasmic poly(A) RNAs and nuclear relocalization of cytoplasmic poly(A) binding protein 1 (PABPC1) in IAV-infected and PA-X overexpressing cells [26,27]. We also demonstrated that upon ectopic overexpression, PA-X can suppress both spliced and unspliced reporter constructs [27]. However, we also showed that in the context of virus infection, PA-X preferentially degrades spliced host transcripts [25]. This indicates that when this potent viral factor is overexpressed, its specificity may be relaxed.

Another IAV host shutoff protein is the nonstructural protein 1 (NS1). Through its N-terminal 80-amino acid double-stranded RNA binding domain (dsRNA), NS1 can interfere with detection of dsRNA by host sensors [28–30], while its C-terminal effector domain is involved in multiple protein-protein interactions [7]. The list of host proteins that can be bound by NS1 is extensive and new interactions continue to be identified [31]. Known as the major viral inhibitor of IFN responses [32,33], NS1 also interferes with general host gene expression by blocking nuclear processing, polyadenylation, and export of mRNAs through binding and inhibition of cleavage and polyadenylation specificity factor 30 (CPSF30) [34,35], nuclear poly(A) binding protein 1 (PABPN1) [36], and nuclear RNA export factor 1 (NXF1) [37,38], respectively. Of these, PABPN1 (also known as PABII) directly affects nascent mRNA poly(A) tail length by bridging the emerging short poly(A) tails and the poly(A) polymerase and stimulating processive poly(A) addition [39]. At some point before or during export or pioneer round of translation, poly(A)-bound PABPN1 is substituted with cytoplasmic PABPC1 [40]. In the nucleus, PABPN1 accumulates in nuclear speckles – subnuclear foci enriched in pre-mRNAs, small nuclear ribonucleoprotein complexes (snRNPs) and serine/arginine rich (SR) proteins involved in splicing [41]. Previous studies have shown that PABPN1 inhibition by NS1 causes shortening of poly(A) tails of nascent host mRNAs and relocalization of PABPN1 from nuclear speckles to a more diffuse distribution throughout the nucleoplasm [36].

The combined effects of NS1 and PA-X in mediating IAV host shutoff have been examined previously [42–45]. These studies suggest that these two proteins co-evolve to ensure an optimal balance between the magnitude of host shutoff and robust viral replication that requires some host gene expression [44]. However, most studies have predominantly focused on the NS1-mediated inhibition of CPSF30, since this function is not conserved in all IAV strains and confers differences in their host shutoff [42,46,47]. In this study, we aimed to determine if the effects of NS1 and PA-X on host gene expression are not simply additive and if there is a functional link between PA-X and NS1 in mediating IAV host shutoff. By using the well characterized laboratory adapted strain A/Puerto Rico/8/1934(H1N1) (PR8) in our model, we examined NS1 effector domain functions independent of CPSF30 inactivation, because PR8 NS1 does not bind CPSF30 [33,47]. To eliminate negative effects of IFN responses on viral protein expression and replication when NS1 is mutated, we conducted most of our analyses in A549 cells lacking MAVS (A549-ΔMAVS, [48]). Our study shows that the NS1 effector domain function is required for PA-X mediated host shutoff and that the NS1-mediated suppression of PABPN1 correlates with nuclear PABPC1 accumulation in IAV-infected cells. We also show that the nuclear relocalization of PABPC1 does not correlate with nuclear poly(A) RNA accumulation in infected cells. This nuclear poly(A) RNA signal accumulation is independent of either PA-X or NS1 functions and is due in part to nuclear retention of viral poly(A) transcripts. Finally, we demonstrate that NS1-mediated host shutoff causes dispersal on nuclear speckles in IAV-infected cells.

## RESULTS

### NS1 effector domain is required for host mRNA depletion and nuclear PABPC1 relocalization in infected cells

To test if nuclear poly(A) RNA and PABPC1 accumulation (previously linked to PA-X activity [26]) was augmented by NS1 effector domain functions, we compared the subcellular distribution of poly(A) RNAs and PABPC1 in A549 cells infected with wild-type (WT) PR8 virus, a PA-X deficient mutant virus (PR8-PA(fs)), an NS1 mutant virus expressing only the N-terminal 80-amino acid RNA-binding domain of NS1 (PR8-NS1(N80)), or a double mutant virus lacking both the PA-X protein and the NS1 effector domain (PR8-PA(fs)-NS1(N80)) (Fig. 1A-C). For this analysis we utilized a combination of immunofluorescence and fluorescence *in situ* hybridization (FISH) microscopy (immunoFISH). This analysis revealed that both the PA-X and the NS1 effector domain were required for nuclear PABPC1 accumulation, which was significantly decreased in cells infected with either PA-X deficient virus, NS1 mutant virus, or the double mutant (Fig. 1A,B). By contrast, the increase in nuclear poly(A) RNA signal did not correlate with nuclear PABPC1 accumulation (Fig. 1A). It also appeared independent of PA-X or NS1 effector domain functions because the significantly increased nuclear to cytoplasmic poly(A) RNA ratio was observed in cells infected with all four recombinant viruses (Fig. 1C). To assess the PA-X mediated host mRNA depletion by the mutant viruses, we isolated total RNA and performed RT-qPCR analysis of the three representative host transcripts ACTB, G6PD, and POLR2A, that we previously reported to be subject to PA-X mediated downregulation [27]. As expected, PA-X deficient virus did not cause significant downregulation of ACTB and G6PD transcripts compared to mock-infected cells (Fig. 1D,E), and the decrease in POLR2A transcript was weaker (Fig. 1F). However, the same phenotype was also observed in cells infected with PR8-NS1(N80) mutant virus that had PA-X gene intact (Fig. 1D-F). These results suggest that NS1 effector domain may be required for PA-X mediated host shutoff. It is possible that in the absence of fully functional NS1, PA-X accumulation is affected because the viral gene expression is inhibited by increased host antiviral responses. Alternatively, the effector domain function may increase PA-X production through general stimulation of viral protein synthesis or specifically through stimulating ribosome frameshifting on PA mRNA. It is also possible that NS1 effector domain can directly stimulate PA-X activity.

**Fig 1.**
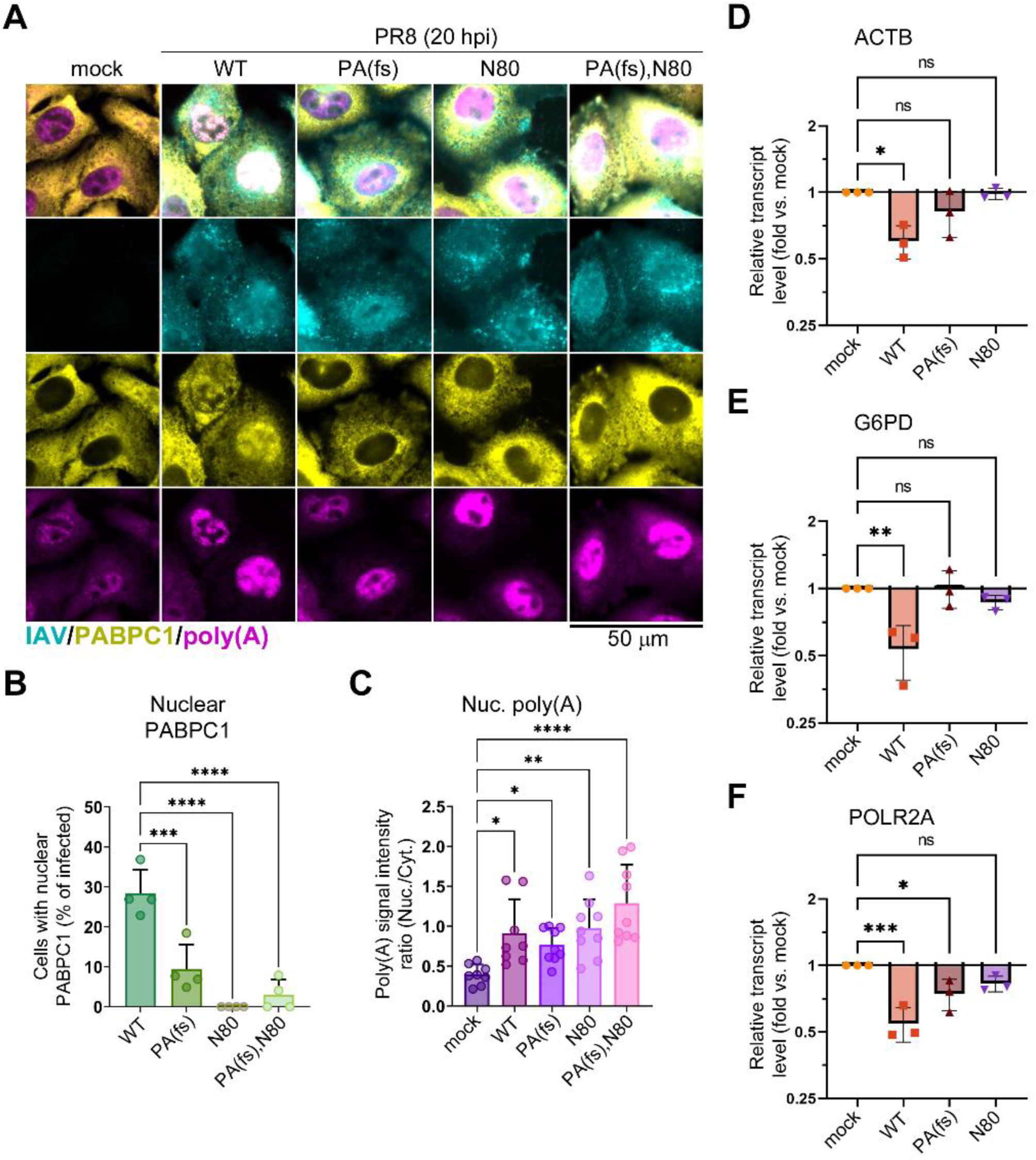
NS1 effector domain is required for PA-X mediated host shutoff and nuclear PABPC1 accumulation in infected cells. A549 cells were mock-infected or infected with the indicated influenza A viruses (PR8 strain) at MOI of 1: wild type (WT), PA(fs) mutant, NS1(N80) mutant (N80), or PA(fs) and NS1(N80) double mutant (PA(fs),N80). (A-C) Cells were fixed and analysed by immunofluorescence microscopy at 20 h post-infection (hpi). (A) Representative immunofluorescence/fluorescence in situ hybridization (ImmunoFISH) microscopy images of cells co-stained using antibodies for influenza A virus structural proteins (IAV, teal), PABPC1 (yellow), and fluorescently labeled oligo(dT) probe (poly(A), magenta). Scale bar = 50 µm. (B) Quantification of mock and PR8-infected cells with nuclear PABPC1 (N = 4). (C) Nuclear to cytoplasmic intensity ratio for poly(A) RNA signal (N = 3), each datapoint represents values obtained from a randomly selected microscopy image containing at least 10 cells. (D, E, F) Total RNA was extracted at 24 hpi and relative levels of ACTB (D), G6PD (E) and POLR2A (F) transcripts were determined by RT-qPCR assay (N = 3). Values were normalized to mitochondrial MT-CYB transcript levels for each sample using ΔΔCt method. (B-F) In all graphs one-way ANOVA and Tukey multiple comparisons tests were done to determine statistical significance (ns: non-significant; ****: p-value < 0.0001; ***: p-value < 0.001; **: p-value < 0.01; *: p-value < 0.05).

### Differences in nuclear PABPC1 accumulation and host mRNA depletion in wild type and NS1 mutant virus infected cells are not due to increased host antiviral response

NS1 activity is crucial for reducing the activation of IFN-mediated responses in infected cells [32], and to regulate both host and viral gene expression [7,49]. Therefore, it is possible that the decrease in PA-X mediated host shutoff by the NS1 effector domain deletion is due to elevated antiviral responses and impaired accumulation of viral proteins, including PA and PA-X. In order to test if IFN responses have major effects on host shutoff phenotypes observed in our experimental system, we employed A549 cells lacking MAVS – the central hub required for IFN induction in virus-infected cells (A549-ΔMAVS) [48]. Indeed, infection of parental A549 cells with PR8-NS1(N80) mutant virus resulted in lower accumulation of PA protein and higher induction of IFN-stimulated genes IFIT1 and ISG15 compared to infection with the WT virus at the same multiplicity of infection (MOI) (Fig. 2A). By contrast, no IFIT1 or ISG15 induction was observed in A549-ΔMAVS cells infected with either the WT or NS1(N80) mutant viruses, and the levels of PA accumulation were more comparable (Fig. 2A). Therefore, we analyzed the subcellular distribution of PABPC1 and poly(A) RNA as well as host transcript depletion in A549-ΔMAVS cells infected with the WT PR8 virus, PR8-PA(fs) mutant virus, and PR8-NS1(N80) mutant virus (Fig. 2B-H). The results were remarkably similar to those obtained in parental A549 cells: only WT PR8 infection resulted in strong nuclear PABPC1 accumulation (Fig. 2B,C), nuclear poly(A) accumulation did not correlate with nuclear PABPC1 and was observed in cells infected by the WT and both mutant viruses (Fig. 2D), and the depletion of ACTB, G6PD, and POLR2A was significantly attenuated by both the PA-X and NS1 mutations (Fig. 2E-G). By contrast, depletion of nuclear non-coding RNA MALAT1 was significantly affected only in NS1(N80) mutant virus infected cells (Fig. 2H). This was consistent with our previous observation that the downregulation of this RNA in PR8-infected cells was PA-X independent [27], and suggests it may be linked to the NS1 effector domain function. Interestingly, when we compared the levels of PA RNA, we saw an approximately 1.5-fold increase in PA levels in PR8-PA(fs) infected cells compared to both the WT PR8 and the PR8-NS1(N80) infected samples (Fig. 2I), indicating that PA-X may affect its own transcript levels. Taken together, results obtained in A549-ΔMAVS cells show that the attenuation of PA-X mediated host shutoff caused by the NS1 effector domain deletion is not due to increased antiviral response and decreased viral replication. They also demonstrate that IFN-mediated antiviral responses in infected cells are not driving an increase in nuclear poly(A) RNA accumulation.

**Fig 2.**
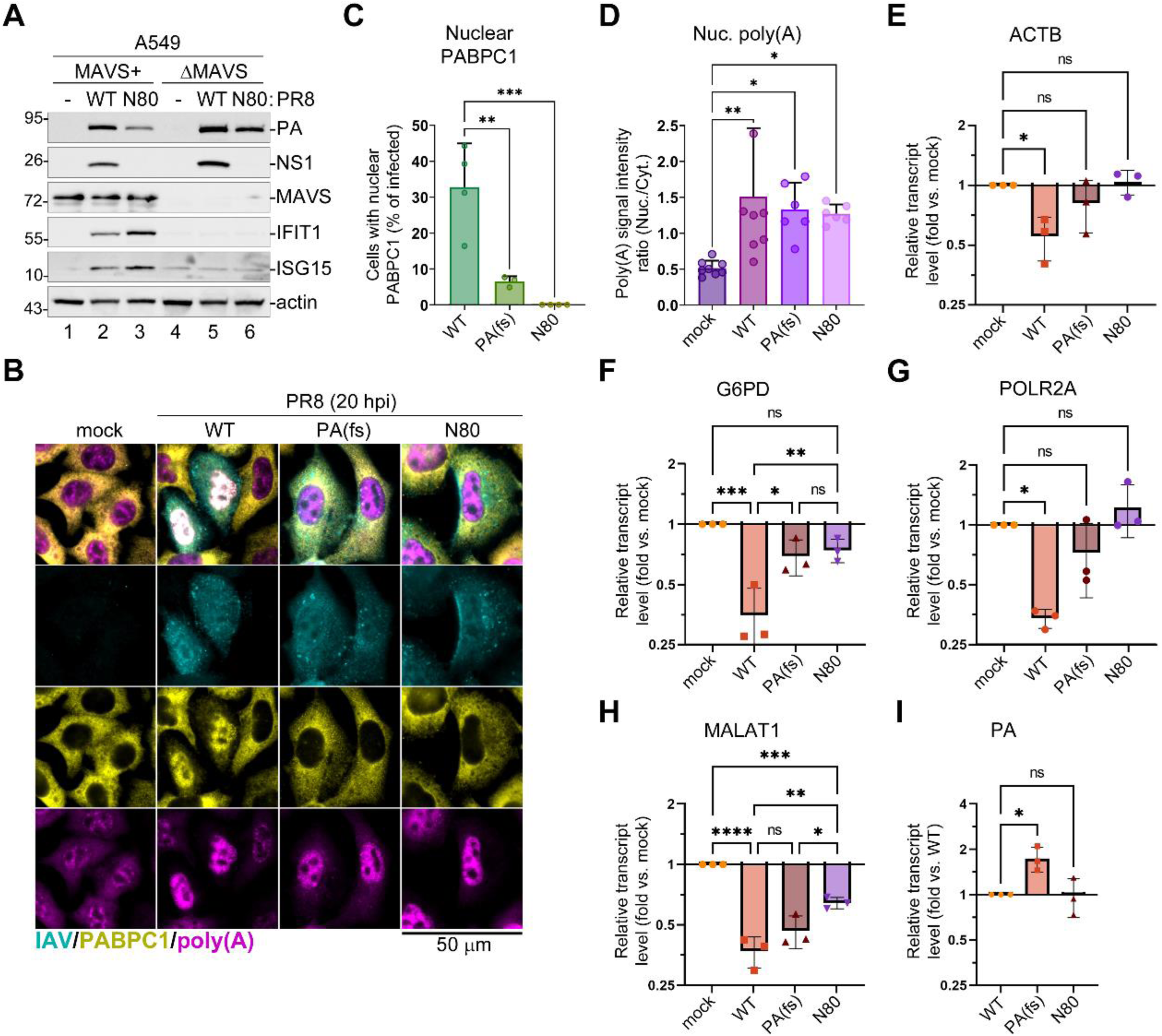
Nuclear poly(A) and PABPC1 accumulation in influenza A virus infected cells are not dependent on MAVS-mediated antiviral response. (A) Parental A549 (MAVS+) and MAVS-deficient (MAVS) cells were mock-infected or infected with either wild-type (WT) or NS1(N80) mutant (N80) PR8 viruses. Levels of the indicated host and viral proteins were analysed using western blotting in whole cell lysates collected at 20 hpi. (B-I) MAVS-deficient A549 cells were mock-infected or infected with the indicated PR8 viruses at MOI of 1: wild type (WT), PA(fs) mutant, NS1(N80) mutant (N80). (B-D) Cells were fixed and analysed by immunofluorescence microscopy at 20 hpi. (B) Representative ImmunoFISH microscopy images of cells co-stained using antibodies for influenza A virus structural proteins (IAV, teal), PABPC1 (yellow), and fluorescently labeled oligo(dT) probe (poly(A), magenta). Scale bar = 50 µm. (C) Quantification of mock and PR8-infected cells with nuclear PABPC1 (N = 4). (D) Nuclear to cytoplasmic intensity ratio for poly(A) RNA signal (N = 3), each datapoint represents values obtained from a randomly selected microscopy image containing at least 10 cells. (E-I) Total RNA was extracted at 24 hpi and relative levels of ACTB (E), G6PD (F), POLR2A (G), MALAT1 (H), and viral PA (I) transcripts were determined by RT-qPCR assay (N = 3). Values were normalized to mitochondrial MT-CYB transcript levels for each sample using ΔΔCt method. (C-I) In all graphs one-way ANOVA and Tukey multiple comparisons tests were done to determine statistical significance (ns: non-significant; ****: p-value < 0.0001; ***: p-value < 0.001; **: p-value < 0.01; *: p-value < 0.05).

### Nuclear poly(A) RNA accumulation is a general phenotype of later stages of influenza A and B virus infection

All Influenza A virus strains are predicted to encode functional PA-X and NS1 proteins. However, the sequences of NS1 and PA-X vary across strains and are subject to adaptive selection [21,44,50]. By contrast, influenza B viruses lack PA-X or a similar host shutoff protein and encode an NS1 with low sequence similarity to the influenza A virus NS1 [21,51]. Therefore, we sought to determine whether nuclear PABPC1 and/or poly(A) RNA accumulation occurs in cells infected with influenza A virus strains other than PR8 or in influenza B virus infected cells. First, we infected A549-ΔMAVS cells with the A/California/7/2009(H1N1) strain of influenza A virus (A/Cal/7) and visualized distribution of PABPC1 and poly(A) RNA using ImmunoFISH at 20 h post-infection (hpi) (Fig. 3A). Compared to PR8 infection (Fig. 2B), A/Cal/7 infection caused even higher fraction of infected cells with nuclear PABPC1 (Fig. 3C). Accumulation of nuclear poly(A) RNA was also evident from microscopy (Fig. 3A) and from measuring the nuclear to cytoplasmic poly(A) RNA signal ratio (Fig. 3D). Second, we infected A549-ΔMAVS cells with the influenza B virus B/Brisbane/60/2008 (B/Bris/60) and subjected them to the same analysis. Consistent with the lack of PA-X host shutoff protein, B/Bris/60 infection did not cause nuclear PABPC1 accumulation (Fig. 3B,C). However, it also resulted in a significant increase in nuclear poly(A) RNA in infected cells (Fig. 3B,D). These results further support the absence of a direct correlation between nuclear PABPC1 and poly(A) RNA accumulation. Nuclear PABPC1 relocalization appears to be characteristic to influenza A viruses that cause PA-X mediated host mRNA depletion, while nuclear poly(A) RNA accumulation also occurs in influenza B virus infected cells and is independent of host shutoff.

**Fig 3.**
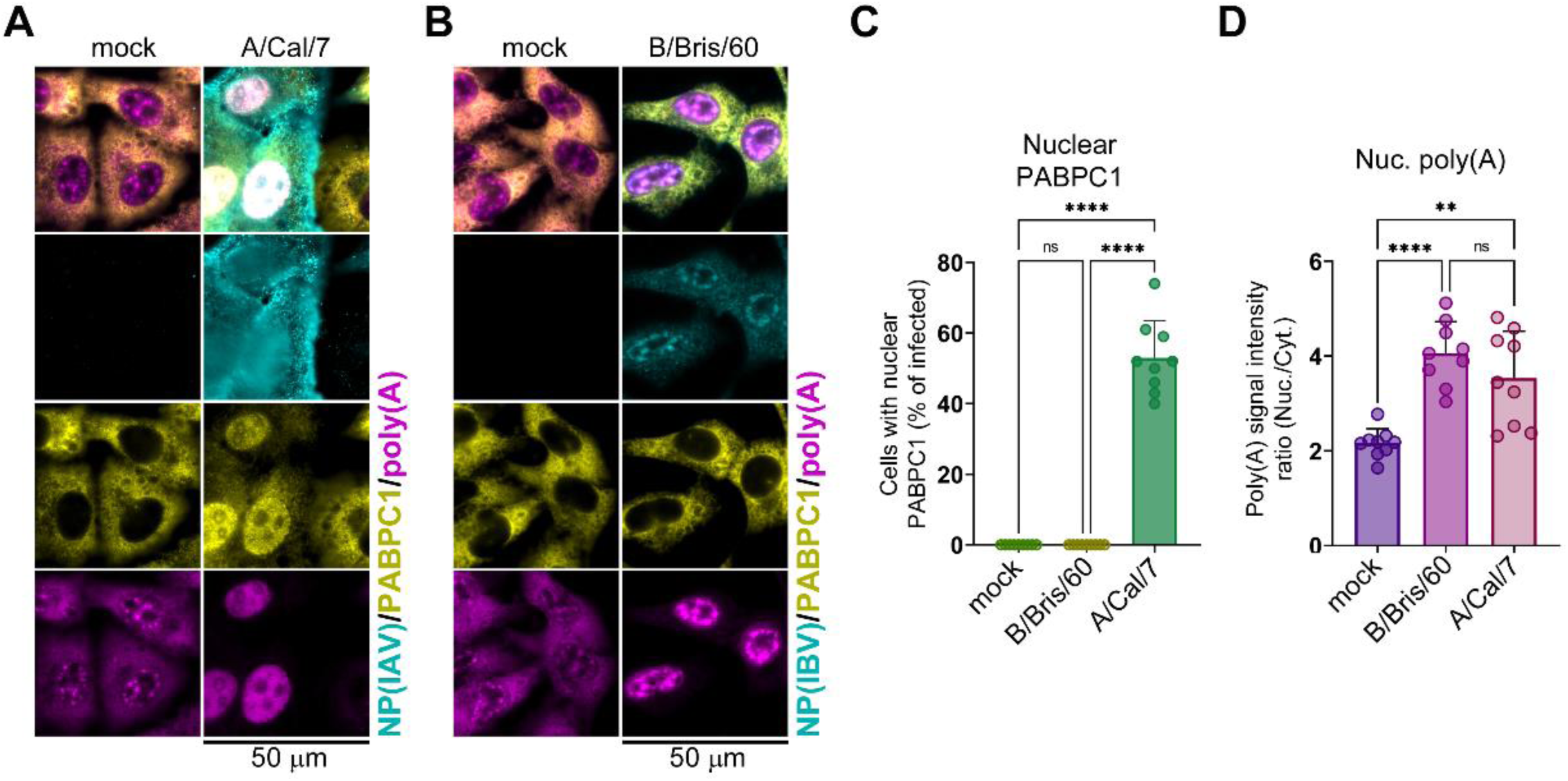
Nuclear poly(A) RNA accumulation in infected cells does not require PA-X activity or nuclear PABPC1 redistribution. (A) MAVS-deficient A549 cells were mock infected or infected with influenza A virus (A/Cal/7) at MOI of 1 and analysed by immunoFISH microscopy at 20 hpi. Cells were co-stained using antibodies for influenza A virus NP (NP(IAV), teal), PABPC1 (yellow), and fluorescently labeled oligo(dT) probe (poly(A), magenta). Scale bar = 50 µm. (B) MAVS-deficient A549 cells were mock infected or infected with influenza B virus (B/Bbris/60) at MOI of 1 and analysed by immunoFISH microscopy at 20 hpi. Cells were co-stained using antibodies for influenza B virus NP (NP(IBV), teal), PABPC1 (yellow), and fluorescently labeled oligo(dT) probe (poly(A), magenta). Scale bar = 50 µm. (C) Quantification of mock-infected, A/Cal/7, and B/Bris/60-infected cells with nuclear PABPC1 (N = 3). (D) Nuclear to cytoplasmic intensity ratio for poly(A) RNA signal (N = 3). In C and D, each datapoint represents values obtained from a randomly selected microscopy image containing at least 20 cells (3 images analyzed per each independent biological replicate). In all graphs one-way ANOVA and Tukey multiple comparisons tests were done to determine statistical significance (ns: non-significant; ****: p-value < 0.0001; **: p-value < 0.01).

### Viral poly(A) RNAs accumulate in the nuclei of influenza A virus infected cells

Nuclear poly(A) RNA accumulation observed in influenza A virus infected cells in our experimental system could correspond to increases in nuclear host mRNAs. However, since we also see this phenotype in PR8-NS1(N80) mutant virus infected cells, this is unlikely to be mediated through an NS1 effector domain-dependent mechanism of mRNA export inhibition [37]. Alternatively, an increase in nuclear poly(A) signal could be due to hyperadenylation of nuclear pre-mRNAs, as has been shown for other viral infections [15], or accumulation of viral poly(A) RNA species. To directly characterise and compare nuclear poly(A) RNA, we isolated nuclear and cytoplasmic RNA fractions from A549-ΔMAVS cells that were mock-infected or infected with the WT PR8 virus, PR8-PA(fs) mutant virus, or PR8-NS1(N80) mutant virus. For fractionation, we used isotonic cytoplasmic RNA extraction buffer containing 0.5% IGEPAL detergent (Fig. 4A-C). To verify that after cytoplasmic lysis the nuclei of infected cells remained intact and contained increased poly(A) RNA, we analyzed cells incubated in buffer without detergent and in full cytoplasmic lysis buffer containing IGEPAL, using smFISH (Fig. 4A). This experiment confirms that lysis efficiently eliminates cytoplasmic poly(A) RNA and cytoplasmic viral genomic RNA signals abundant in the control cells, while preserving increased poly(A) signal in the nuclei of infected cells (Fig. 4A). A total of six independent biological replicates of nuclear and cytoplasmic RNA isolation were performed. On average, 2-3 times more RNA was isolated from the cytoplasm compared to nuclear fractions (S1A Fig). All individual RNA preparations were combined in pooled fractions. Further analysis confirmed that the isolated RNA fractions contained intact 28S and 18S ribosomal RNAs (Fig. 4B), and that the nuclear fractions were substantially enriched in NEAT1 transcript that normally accumulates in nuclear paraspeckles [52] (Fig. 4C). Subsequently, poly(A) RNA was isolated from all nuclear fractions and the cytoplasmic fractions from mock and WT PR8 infected cells and analysed by direct RNA sequencing using Oxford Nanopore long-read sequencing (S1 Table and S1B Fig). First, we compared nuclear and cytoplasmic poly(A) RNAs between mock and the WT PR8 infected samples (Fig. 5). Consistent with our previous analysis using RNAseq [25], in infected cells approximately half of total nuclear and cytoplasmic poly(A) reads mapped to viral genes. At the same time, the nuclear fraction contained more viral than host reads while more host reads were present in the cytoplasm (Fig. 5A). Of the viral poly(A) reads, the NP transcript was the most abundant in both the nucleus and cytoplasm, making it the most abundant single mRNA species in the infected cell (Fig. 5B and S2 Table). The least abundant viral transcript was PA, followed by NEP. The overall relative abundance of viral transcripts was similar between the nucleus and the cytoplasm, with exception of M2 mRNA which was more abundant in the cytoplasmic fraction (5% of all viral reads) than in the nucleus (2%), and the PB2 mRNA, which was more abundant in the nuclear fraction (8% of viral reads) than in the cytoplasm (4%) (Fig. 5B). A total of 1,249 host transcripts were identified in all mock and virus-infected poly(A) RNA preparations (Fig. 5C). Significantly more host transcripts (1,898) were only found in mock-infected cells. These were predominantly corresponding to lower abundance RNAs with fewer than 150 reads that were likely eliminated due to a combination of virus-induced host shutoff and the influx of viral transcripts. Very few host transcripts were only detected in virus-infected samples (14 in the nucleus, 35 in the cytoplasm, and 20 in both fractions). Consistent with our previously reported preferential targeting of multiply spliced host mRNAs by the PA-X host shutoff, this group was enriched in intron-less transcripts: processed pseudogenes of ribosomal proteins, small non-coding RNAs (small nucleolar RNAs, signal recognition particle RNAs, 5.8S ribosomal RNAs), and several stress response genes that escaped PA-X mediated degradation and/or were induced in response to infection (S1C Fig). By contrast, of the 100 most abundant host poly(A) transcripts (ranked by the combined read number in two mock samples), all were decreased in the cytoplasmic fraction and majority were decreased in the nuclear fraction of PR8-infected cells (Fig. 5D). Of the few nuclear RNAs that were not downregulated, FTL and RPL39 were the most increased (1.6 fold). Overall, FTL was the most abundant host poly(A) transcript in both mock and virus-infected cells, which is consistent with our previous RNAseq analysis of the transcriptome of this cell type [25]. However, in the nucleus of infected cells the read count of the single viral NP transcript was 40 times higher than this most abundant host transcript, illustrating that viral and not host mRNAs were likely responsible for overall increased nuclear poly(A) signal. Next, we assessed the poly(A) tail lengths of host transcripts using the Nanopolish-polyA algorithm [53]. This analysis revealed that the average poly(A) length significantly decreased in infected cells compared to mock infected cells, indicating that hyperadenylation of host transcripts was not responsible for increased poly(A) signal in the nucleus (Fig. 5E). To illustrate changes in polyadenylation of individual host transcripts we selected ACTB and GAPDH that are often used as loading controls on western and northern blots. While polyadenylation of ACTB did not change significantly (Fig. 5F), the average poly(A) length of GAPDH transcripts decreased from 83 to 66 nt in the nucleus and from 73 to 56 nt in the cytoplasm (Fig. 5G). As a control for poly(A) tail length estimation, we used *Saccharomyces cerevisiae* ENO2 spike-in RNA which has a defined poly(A) tail length of 30 nt. Nanopolish-poly(A) overestimated the ENO2 poly(A) tail length to be between 39 and 40 nt but this length remained consistent with less than 1 nt or 2.5% difference between samples (Fig. 5H) and the overall distribution of ENO2 poly(A) tail lengths was the same as was reported previously using this algorithm [53]. Given that we observed much larger differences (20-30%) in host transcript poly(A) tail length, this method was suitable for relative poly(A) length estimation but did not produce the most accurate absolute length measurements.

**Fig 4.**
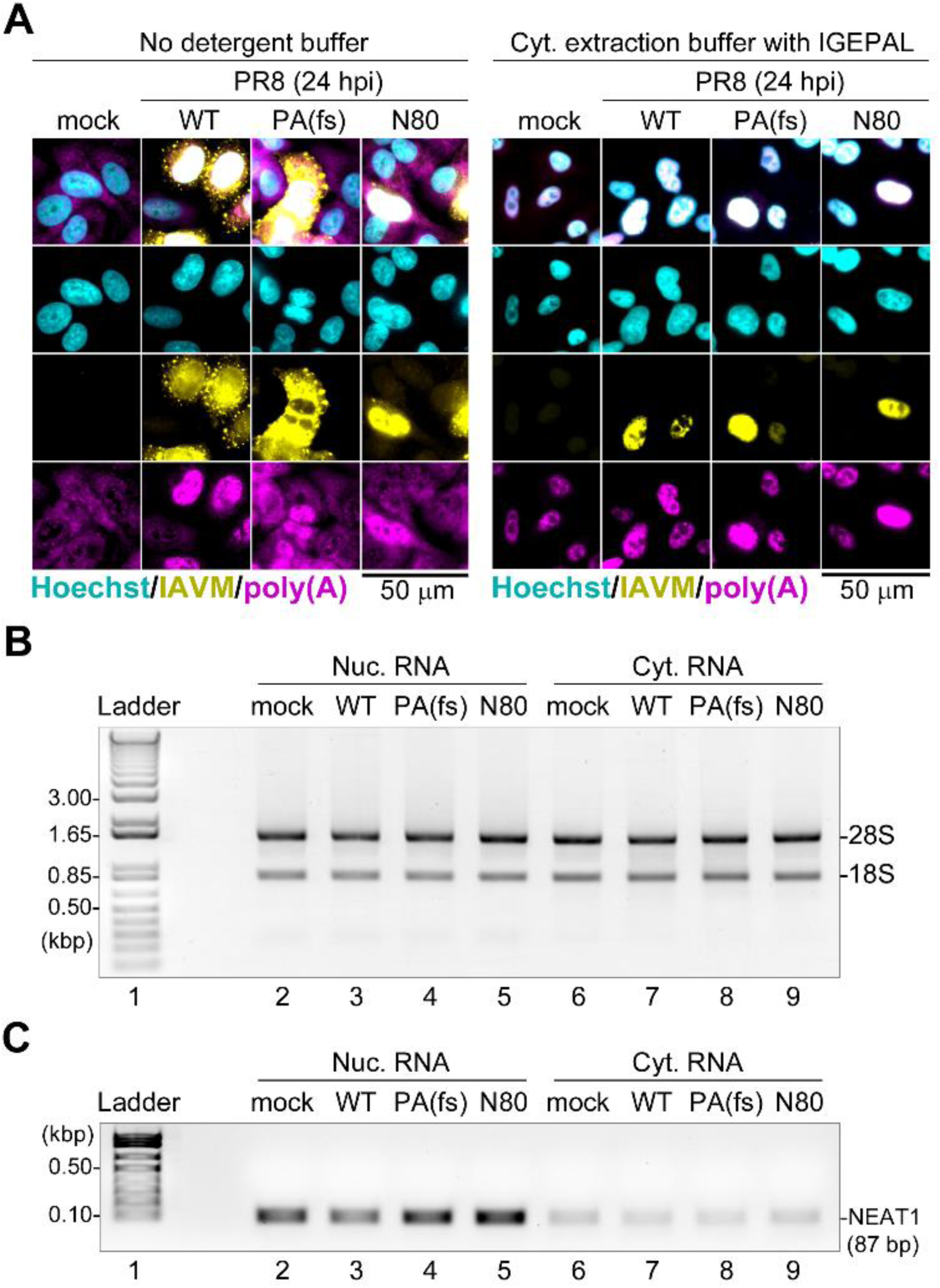
Nuclear and cytoplasmic RNA isolation. (A) Fluorescence microscopy images of MAVS-deficient A549 cells mock-infected or infected with the indicated PR8 viruses at MOI of 1: wild type (WT), PA(fs) mutant, NS1(N80) mutant (N80). At 24 hpi, cells were incubated in the cytoplasmic extraction buffer with (right) or without (left) IGEPAL detergent prior to fixation and smFISH analysis. Infected cells were visualised using smFISH probe set for viral genomic segment 7 (IAVM, yellow) and total poly(A) RNA was visualized using oligo(dT) probe (poly(A), magenta). Cell nuclei were stained with Hoechst dye (teal). Scale bar = 50 µm. (B) Agarose gel analysis of total nuclear (Nuc.) and cytoplasmic (Cyt.) RNA fractions obtained from the indicated mock-infected and PR8-infected cells. 1% agarose “bleach gel” with ethidium bromide staining was used as described in (REF), the RNA fluorescence image was inverted for the panel presentation. Each lane contains a pooled sample from six independent replicates. Positions of the 28S and 18S ribosomal RNA bands are indicated. (C) Agarose gel analysis of NEAT1 RNA amplicons obtained using semi-quantitative PCR. The template cDNAs were obtained from the indicated nuclear (Nuc.) and cytoplasmic (Cyt.) RNAs corresponding to those shown in panel B. bp: base pairs.

**Fig 5.**
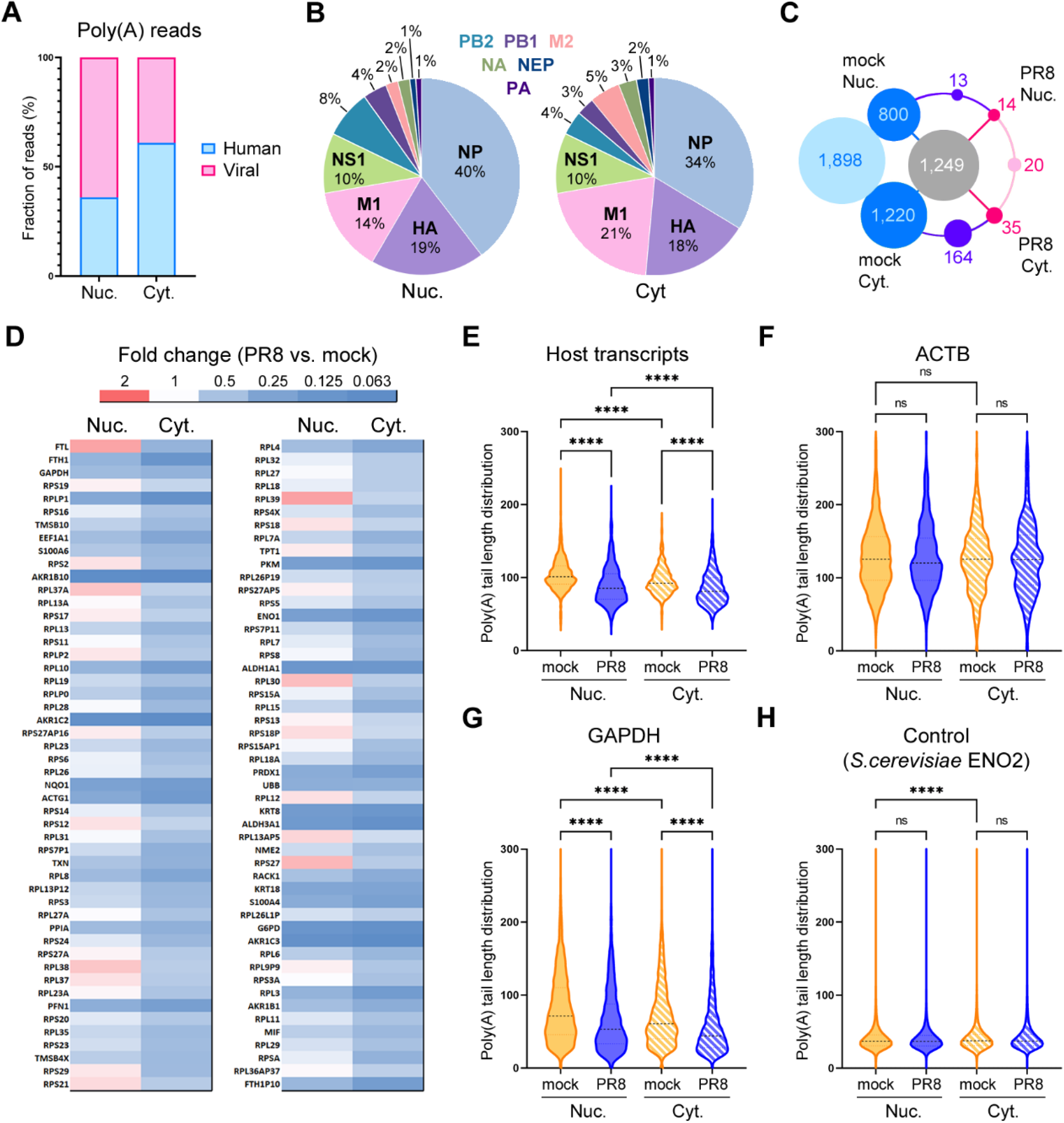
Viral poly(A) transcripts accumulate in the nucleus of infected cells. Analysis of nuclear (Nuc.) and cytoplasmic (Cyt.) poly(A) RNAs isolated at 24 hpi from mock infected and influenza A virus infected A549-ΔMAVS cells using Oxford Nanopore direct RNA sequencing. (A) Proportion of influenza A virus (Viral, pink) and host cell (Human, blue) poly(A) reads. (B) Relative abundances of each of 10 major viral mRNA transcripts plotted as percent of total viral reads in the nuclear (left) and cytoplasmic (right) RNA fractions. (C) Number of shared and unique host transcripts identified in each RNA sample represented as a wheel diagram. Number of transcripts identified in all four samples is shown in grey circle in the center, identified only in mock-infected RNA samples in blue, and identified only in virus-infected RNA samples (PR8) in pink. Numbers of exclusively cytoplasmic and exclusively nuclear transcripts common for both mock and PR8 samples are shown in purple. (D) Heat map showing the relative change in the levels of the 100 most abundant host poly(A) transcripts in influenza A virus-infected cells compared to mock-infected cells. (E) Violin plot showing the distribution of host transcript poly(A) tail lengths as determined using Nanopolish-poly(A) in mock and influenza A virus-infected (PR8) nuclear and cytoplasmic RNAs. (F-H) Distribution of individual read poly(A) tail lengths for the indicated representative transcripts: (F) human ACTB; (G) human GAPDH; (H) S. cerevisiae ENO2 spike-in control mRNA. (E-H) On all plots, one-way ANOVA and Tukey’s multiple comparisons test was done to determine statistical significance (ns: non-significant, ****: p-value < 0.0001).

### NS1 effector domain functions have a major effect on nuclear poly(A) RNA composition in infected cells

To further examine the impacts of PA-X and NS1 host shutoff on polyadenylation and accumulation of host and viral transcripts in the nuclei of infected cells, we compared nuclear poly(A) RNA between samples isolated from A549-ΔMAVS cells infected with the WT PR8, PA(fs) mutant virus, and the NS1(N80) mutant virus (Fig. 6). In both mutant virus infection samples, viral transcripts constituted a lower share of reads compared to WT (Fig. 6A). This could be due to direct effects of host shutoff on viral mRNA export. Nevertheless, the relative abundances of viral transcripts were similar in all three samples, with NP being the most abundant viral nuclear mRNA and the single most abundant nuclear transcript in infected cells (Fig. 6B). Of the 10 major influenza mRNAs, the alternatively spliced M1/M2 and NS1/NEP transcripts were most affected by the NS1 effector domain deletion: relative abundance of unspliced M1 and NS1 transcripts decreased 2 and 5 fold, respectively, while relative abundance of spliced transcripts increased (1.5 fold for M1 and 2 fold for NEP) (Fig. 6B). Among host nuclear transcripts, the general trend in their change was towards higher abundance in mutant virus infected cells compared to WT (Fig. 6C). Transcripts that were most depleted in the nuclei of the WT virus infected cells (e.g. AKR1B10, ALDH3A1) were decreased to a lesser degree, while transcripts that increased in abundance (e.g. FTL, RPL30) increased even more (Fig. 6C). This was not surprising, considering the lower relative abundance of viral transcripts, especially in PR8-NS1(N80) infected cells (Fig. 6A). Next, we estimated the average poly(A) tail length of host nuclear transcripts (Fig. 6D-F). In the WT and the PA(fs) mutant virus infected cells, the average poly(A) tail length decreased to a similar extent compared to mock infected cells. By contrast, poly(A) tail length slightly increased compared to mock in NS1(N80) mutant virus infected cells (Fig. 6D-F). Interestingly, we also observed significant lengthening of the average poly(A) tails of the viral NP and most other viral transcripts in PA(fs) mutant virus infected cells (from 85 to 92 nt), and in NS1(N80) mutant virus infected cells (from 85 to 111 nt) (Fig. 6G and S2 Table). Taken together, these results indicate that influenza A virus host shutoff leads to a general decrease in poly(A) tails of both host and viral transcripts, with NS1 effector domain playing a major role in this phenotype. For the general increase in poly(A) RNA signal observed in the nuclei of the NS1(N80) mutant virus infected cells, the lower influx or retention of viral transcripts is compensated by the impaired downregulation of host transcripts and the general increase in poly(A) lengths.

**Fig 6.**
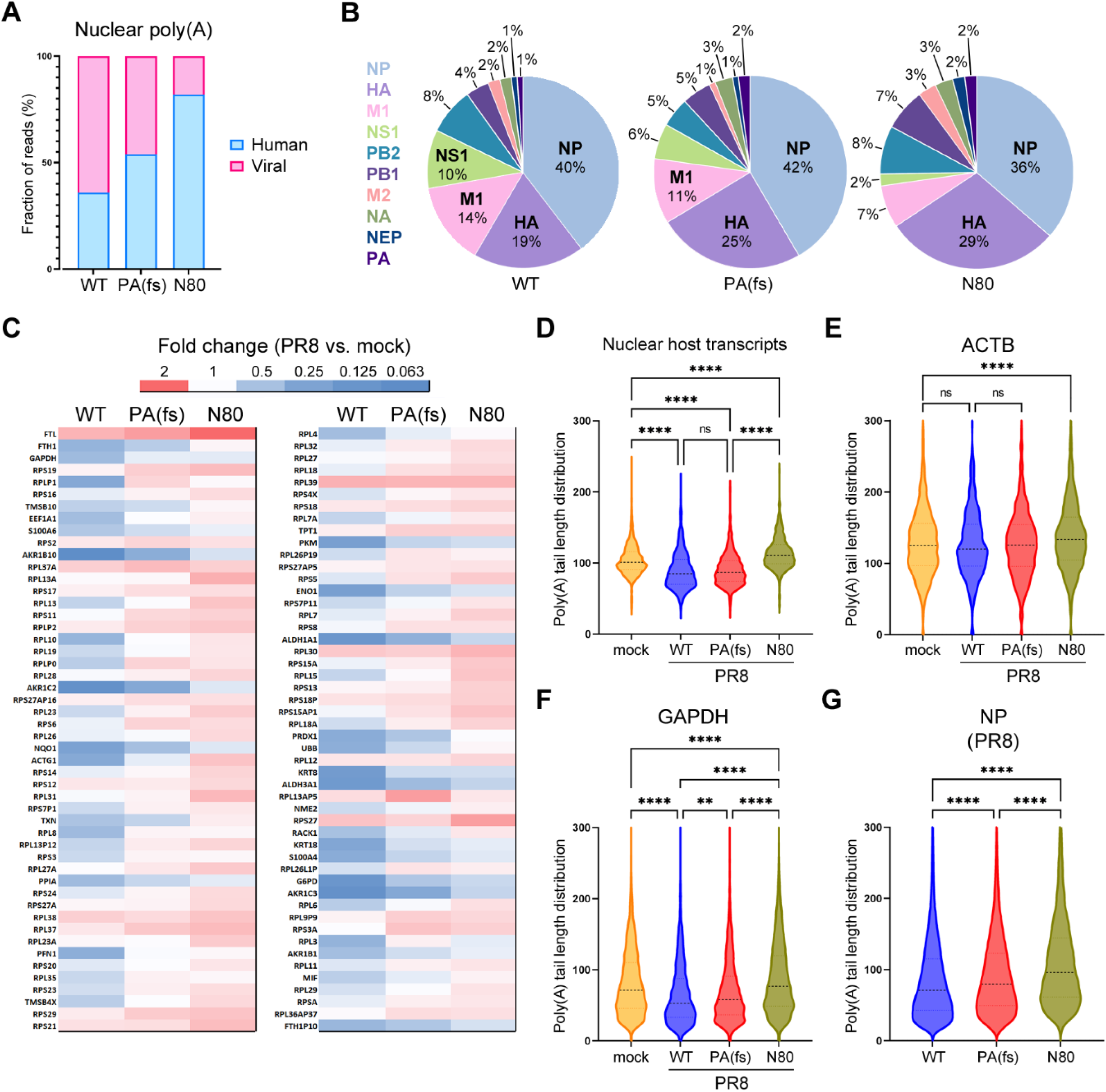
NS1 effector domain function is responsible for general decrease in poly(A) tail lengths of nuclear transcripts. MAVS-deficient A549 cells were mock-infected or infected with the indicated PR8 viruses at MOI of 1: wild type (WT), PA(fs) mutant, or NS1(N80) mutant (N80). Nuclear poly(A) RNAs were isolated at 24 hpi and analyzed using Oxford Nanopore direct RNA sequencing. (A) Proportion of influenza A virus (Viral, pink) and host cell (Human, blue) poly(A) reads. (B) Relative abundances of each of 10 major viral mRNA transcripts plotted as percent of total viral reads in the nuclear RNA fractions. (C) Heat map showing the relative change in the levels of the 100 most abundant host poly(A) transcripts in the nuclear fractions of cells infected with the wild-type and the indicated mutant viruses compared to mock-infected cells. (D) Violin plot showing the distribution of nuclear host transcript poly(A) tail lengths as determined using Nanopolish-poly(A). (E-G) Distribution of individual read poly(A) tail lengths for the indicated representative transcripts: (E) human ACTB; (E) human GAPDH; (H) viral NP mRNA. (A-G) On all panels except panel G, the data for nuclear mock and wild-type PR8 infected cell RNA analysis is duplicated from figure 5 to allow direct comparison with other conditions. On all plots, one-way ANOVA and Tukey’s multiple comparisons test was done to determine statistical significance (ns: non-significant, ****: p-value < 0.0001; **: p-value < 0.01).

### NS1 sequesters PABPN1 away from nuclear speckles

Having demonstrated that the NS1 effector domain deletion impairs influenza A virus host shutoff and abolishes nuclear PABPC1 accumulation, we examined whether similar effects could be observed using point mutations in the NS1 effector domain. To this end, we generated recombinant PR8 viruses with W187R substitution in the NS1 that inhibits effector domain dimerization [54], and the double alanine substitution of adjacent highly conserved surface exposed amino acids I123 and M124, originally reported to be involved in viral mRNA synthesis regulation and, in addition, dsRNA activated protein kinase (PKR) binding and inhibition [55]. While the W187R substitution in NS1 had no effect on nuclear PABPC1 accumulation, which was comparable to WT PR8 virus infection, NS1(123A,124A) mutant virus did not cause this phenotype (Fig. 7A,B). In this respect, the double amino acid substitution phenocopied the complete NS1 effector domain deletion in PR8-NS1(N80) virus (Fig. 2B,C). Therefore, we included PR8-NS1(123A,124A) virus in our next series of experiments. One of the NS1 protein interactors in the nuclei of infected cells is PABPN1, and the NS1 effector domain is required for this interaction and interference with PABPN1 function and localization to nuclear speckles [36]. We analyzed the subcellular distribution of PABPN1 in mock infected cells and in WT or mutant PR8 virus infected cells and observed a similar phenotype first reported by Chen et al. [36]: in uninfected cells PABPN1 concentrated in nuclear speckles and in WT PR8 virus infected cells PABPN1 signal was more diffusely distributed and decreased in intensity (Fig. 7C). The same phenotype was observed in cells infected with PR8-PA(fs) virus with an intact NS1 gene. By contrast, in cells infected with either NS1(N80) or NS1(123,124A) mutant viruses, the PABPN1 staining pattern was similar to uninfected cells (Fig. 7C). Because the immunofluorescence signal for PABPN1 was weaker in WT virus infected cells, we wanted to see if the virus downregulated PABPN1 expression using western blotting. We did not detect significant downregulation of PABPN1 (Fig. 7D,E). Taken together, these results demonstrate that the poly(A) tail shortening correlates with NS1-mediated PABPN1 sequestration away from the nuclear speckles observed in WT and PA(fs) mutant virus infected cells, while nuclear PABPC1 accumulation only partially correlates with this NS1-dependent phenotype and still requires PA-X.

**Fig7.**
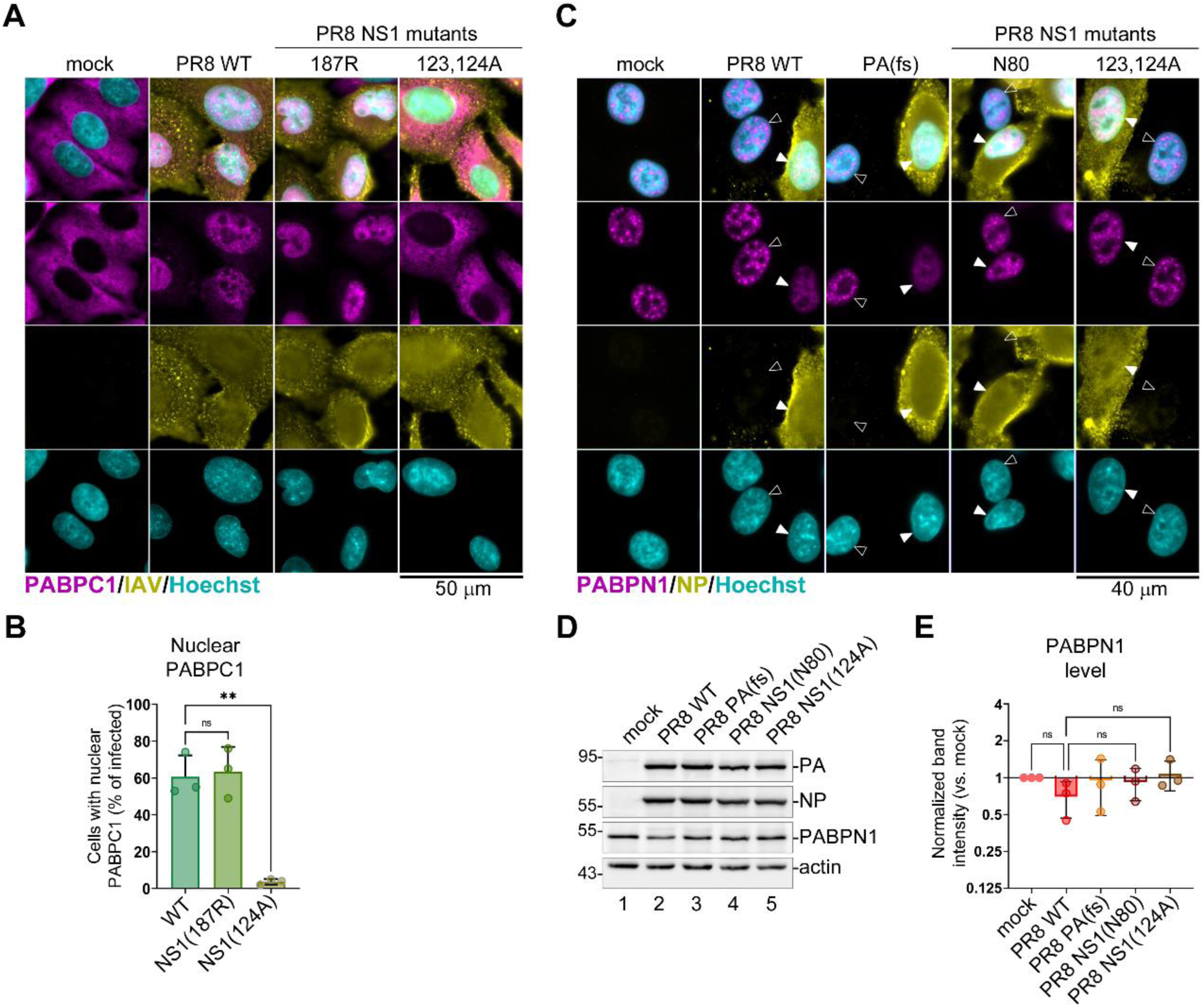
NS1 disrupts PABPN1 localization to nuclear speckles. (A-C) Fluorescence microscopy analyzes of MAVS-deficient A549 cells mock-infected or infected with the indicated PR8 viruses at MOI of 1 at 24 hpi: wild type (WT), PA(fs) mutant, NS1(W187R) mutant (187R), or NS1(I123A,M124A) mutant (124A). (A) Representative immunofluorescence microscopy images of cells co-stained using antibodies for influenza A virus structural proteins (IAV, yellow) and PABPC1 (magenta). Nuclei were stained with Hoechst dye (teal). Scale bar = 50 µm. (B) Quantification of infected cells with nuclear PABPC1 performed on immunostained cells represented in panel A (N = 3). (C) Representative immunofluorescence microscopy images of cells co-stained using antibodies for influenza A virus nucleoprotein (NP, yellow) and PABPN1 (magenta). Nuclei were stained with Hoechst dye (teal). Filled arrowheads highlight nuclei of infected cells, open arrowheads highlight bystander uninfected cells. Scale bar = 40 µm. (D) Levels of the indicated host and viral proteins were analysed using western blotting in whole cell lysates collected at 24 hpi. (E) Relative intensity of PABPN1 band was quantified from western blot analyses represented in panel D. In each replicate, values were normalized to actin (N = 3). (B,E) One-way ANOVA and Tukey’s multiple comparisons test was done to determine statistical significance (ns: non-significant; **: p-value < 0.01).

### Influenza A virus causes dispersal of nuclear speckles in infected cells

We observed redistribution of PABPN1 from nuclear speckles to a more diffuse pattern in the nuclei of virus-infected cells (Fig. 7C). Next, we wanted to test if nuclear speckles themselves were altered by influenza A virus. To visualize nuclear speckles by confocal microscopy, we used smFISH probe set for MALAT1 non-coding RNA (Fig. 8A) and the polyclonal antibody to SR proteins (Fig. 8B), which are components of nuclear speckles. Compared to bystander uninfected cells that had well defined nuclear MALAT1 staining with speckled pattern, WT PR8 virus infected cells had little to no MALAT1 staining (Fig. 8A). In addition, the SR staining showing co-localization with PABPN1 in defined nuclear foci in uninfected cells, was more dispersed in smaller more numerous foci in infected cell nuclei (Fig. 8B). These results demonstrate that influenza A virus infection disperses nuclear speckles.

**Fig 8.**
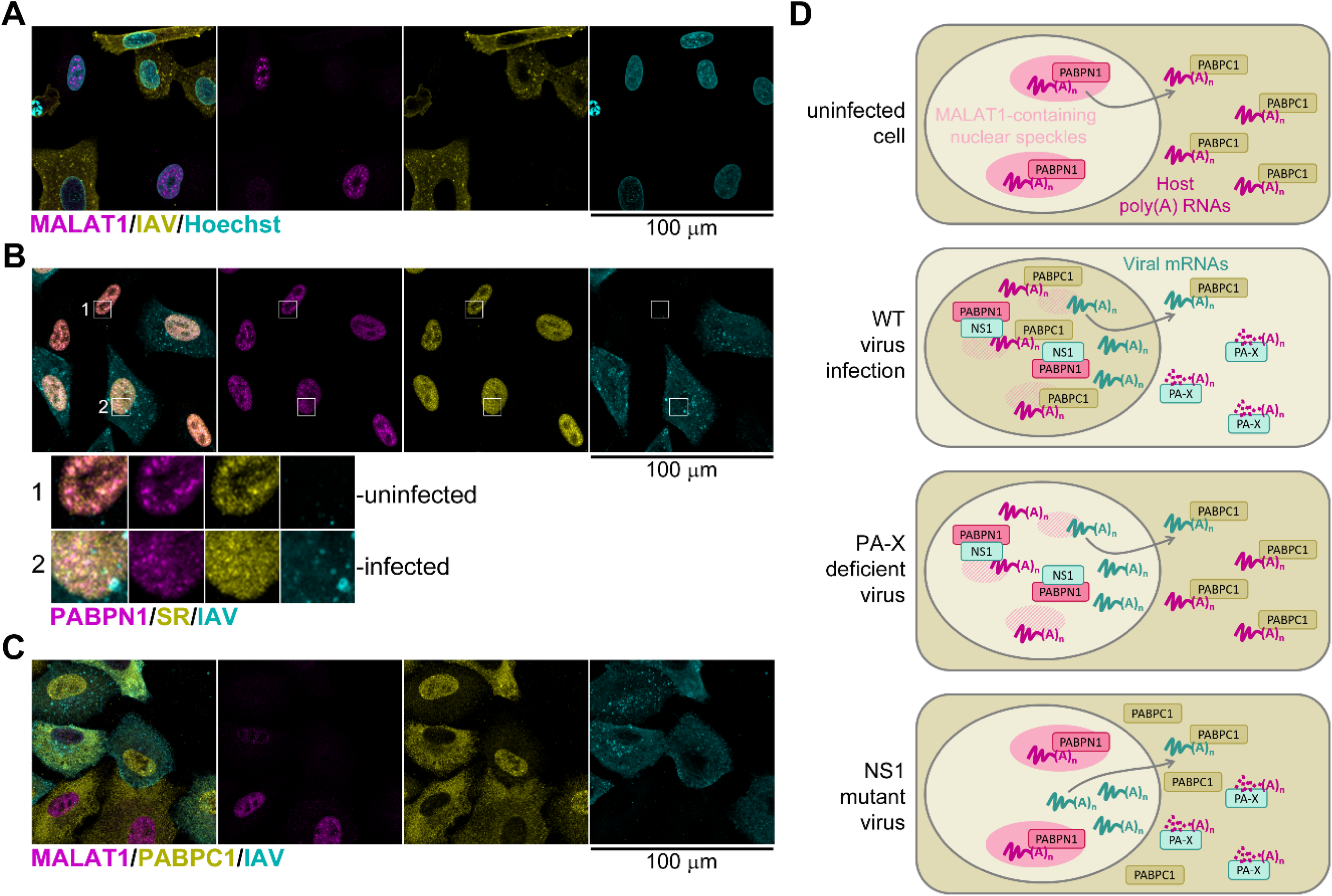
Influenza A virus host shutoff causes dispersal of nuclear speckles in infected cells. (A-C) Confocal fluorescence microscopy analyzes of A549-ΔMAVS cells infected with influenza A virus (PR8 strain) at MOI of 1 at 24 hpi. Scale bars = 100 µm. (A) Representative immunoFISH microscopy image of cells co-stained using antibody for influenza A virus nucleoprotein (NP, yellow) and the smFISH probe set for the nuclear MALAT1 transcript (magenta). Nuclei were stained with Hoechst dye (teal). (B) Immunofluorescence microscopy image of cells co-stained using antibodies for influenza A virus nucleoprotein (NP, teal), PABPN1 (magenta), and SR proteins (yellow). Outsets show enlarged regions of the nuclei of a representative uninfected bystander cell (1) and virus-infected cell (2). (C) Representative immunoFISH microscopy image of cells co-stained using antibody for influenza A virus nucleoprotein (NP, teal), PABPC1 (yellow), and the smFISH probe set for the nuclear MALAT1 transcript (magenta). (D) Working model for the concerted action of NS1 and PA-X proteins in mediating nuclear accumulation of PABPC1. In uninfected cells (top diagram), nascent transcripts traffic through nuclear speckles containing MALAT1 RNA and PABPN1 protein. Upon cytoplasmic export, host mRNAs associate with PABPC1 that enhances their translation. In infected cells (upper middle diagram), PA-X depletes host poly(A) RNAs, causing excess of free PABPC1 with unmasked nuclear localization signal. Simultaneously, NS1 protein interferes with the processing and maturation of host pre-mRNAs, in part through sequestering PABPN1 protein. This results in dispersal of nuclear speckles and depletion of MALAT1 RNA. Sequestration of PABPN1 causes accumulation of nuclear PABPC1 that can preferentially bind nuclear poly(A) RNAs. When PA-X production is inhibited by frameshift site alteration in PA(fs) mutant virus (lower middle diagram), nuclear import of PABPC1 is diminished because the host cytoplasmic mRNAs are not sufficiently depleted. When NS1-mediated sequestration of PABPN1 is disrupted by mutations (bottom diagram), nuclear accumulation of PABPC1 is blocked by poly(A)-bound PABPN1.

## DISCUSSION

The discovery of the endonuclease PA-X over a decade ago filled an important gap in our understanding of the mechanisms of host shutoff by IAV [18]. Our previous functional analyses identified nuclear accumulation of PABPC1 as a hallmark of PA-X mediated host shutoff in both infected cells and cells ectopically overexpressing PA-X protein [26,27]. Accordingly, we proposed a model in which the depletion of cytoplasmic mRNAs by PA-X results in an excess of PABPC1 protein not bound to poly(A) tails, which unmasks nuclear localization signal of PABPC1 located in the RNA-binding domain [19]. This model was consistent with the reported mechanism governing nucleocytoplasmic shuttling of PABPC1 [56–58]. It was also informed by our analysis of the SARS-CoV2 host shutoff protein Nsp1 that causes similar nuclear PABPC1 relocalization [59], and a previously proposed model for nuclear PABPC1 accumulation caused by KSHV host shutoff nuclease SOX and its homologue from the murine gammaherpesvirus 68 (MHV68) called muSOX [15]. That work by the Glaunsinger group and their later studies also described how aberrant nuclear accumulation of PABPC1 caused hyperadenylation of mRNAs [15,56]. In our IAV infection model we observed a striking increase in nuclear poly(A) signal in virus-infected cells, suggesting that a similar hyperadenylation phenotype could result from nuclear PABPC1 accumulation [26].

In this study, for the first time, we provide evidence that NS1 has a major effect on PA-X activity and is required for nuclear PABPC1 accumulation in IAV infected cells. In addition to inhibiting host pre-mRNA processing, polyadenylation, and export, NS1 regulates viral gene expression by affecting alternative splicing of segment 7 transcript [60], viral mRNA export [49], and translation [61,62]. Consequently, besides contributing to general host shutoff, NS1 can influence PA-X function specifically by promoting its synthesis or altering the subcellular distribution of its target mRNAs. In this work, we show that IAV infection dramatically alters nuclear organization by dispersing nuclear speckles and depleting its major RNA constituent MALAT1 (a.k.a. NEAT2, [63]) in a PA-X independent manner. In addition, the NS1 effector domain mediated PABPN1 sequestration correlates with nuclear accumulation of PABPC1. This function of NS1 requires interactions mediated by amino acids I123 and M124, but is independent of effector domain dimerization. In infected cells, nuclear PABPC1 accumulation is the result of a concerted action by both PA-X and NS1. We propose a working model for nuclear PABPC1 relocalization in IAV-infected cells, in which NS1 alleviates competition from PABPN1 for binding nuclear poly(A) RNAs, allowing PABPC1 to accumulate in the nucleus following cytoplasmic poly(A) RNA depletion by PA-X (Fig. 8D). By contrast, nuclear poly(A) accumulation in IAV-infected cells detected using immunoFISH is independent of either PA-X, NS1 effector domain, or nuclear PABPC1. Notably, we observe similar nuclear poly(A) RNA signal accumulation in cells infected with influenza B virus that does not encode PA-X and does not cause nuclear PABPC1 relocalization.

To characterize polyadenylated transcripts that accumulate in the nuclei of IAV-infected cells, we isolated poly(A) RNAs from the nuclear and the cytoplasmic fractions and analysed them using nanopore sequencing. Our analysis revealed that the large proportion of nuclear poly(A) RNAs are viral, and that the NP mRNA is the single most abundant poly(A) transcript in infected cells. Overall, decrease in host poly(A) RNAs was stronger in the cytoplasmic fraction, with 100 of the most abundant mRNAs all being downregulated compared to mock-infected cells. In the nucleus, while some of these transcripts increased in abundance relative to uninfected cells, their levels were still much lower than those of the viral mRNAs. To assess poly(A) tail lengths of isolated RNAs we employed Nanopolish-polyA analysis [53]. As reported previously [53,64], in uninfected cells the average poly(A) tail length was shorter in the cytoplasmic fraction compared to the nuclear fraction. In WT virus infected cells, we did not observe hyperadenylation of host transcripts, and instead the average poly(A) tail length decreased compared to mock-infected cells. This observation is consistent with NS1-mediated inhibition of PABPN1 function. Another interesting phenotype that may be directly linked to NS1-mediated PABPN1 inhibition is the detection of polyadenylated small nucleolar RNAs (snoRNAs) SNORD3B-2 and SNHG25 in WT infected cells and polyadenylated 5.8S ribosomal RNAs and signal recognition particle RNAs in the nuclear fraction of both WT and PA(fs) mutant virus-infected cells, but not in mock-infected or NS1(N80) mutant virus infected cells. Normally these nuclear RNAs are not polyadenylated [65], however a study by Lemay *et al.* showed that deletion of the PABPN1 homolog in fission yeast called Pab2 leads to accumulation of polyadenylated snoRNAs [66]. The mechanism of accumulation is linked to impaired PABPN1-mediated recruitment of nuclear exosome that trims transiently added poly(A) tails of these small RNAs as part of their maturation pathway [66,67]. In the future, it will be interesting to test if similar mechanism of aberrant small nuclear RNA polyadenylation is caused by NS1-mediated PABPN1 inhibition and what effect it has on viral replication or host shutoff. As for the mRNAs, we observed a small but significant increase in average poly(A) tail length of host and viral nuclear transcripts in NS1(N80) mutant virus infected cells. However, this hyperadenylation was not driven by nuclear PABPC1 which does not occur in PR8-NS1(N80) mutant virus infection. Unlike NS1 effector domain deletion, attenuation of PA-X production by PA(fs) mutation had no significant effect on host nuclear poly(A) RNA tail lengths.

Overall, our study demonstrates that even in the absence of CPSF30 binding, NS1 has major effect on IAV host shutoff by inhibiting PABPN1 and enhancing PA-X activity. Even in the absence of MAVS-dependent antiviral signaling, PA-X mediated host mRNA degradation was impaired by the deletion of the NS1 effector domain. Future studies should directly address the question of whether NS1 is required for efficient PA-X synthesis in infected cells. Unfortunately, this seemingly trivial task will require new method development and/or reagent development for PA-X detection. We have tested a number of commercially available PA-X antibodies and were unable to confirm that they reliably detect PA-X even upon ectopic overexpression. Our analysis of PA RNA and protein expression shows that in PR8-NS1(N80) mutant virus infected A549-ΔMAVS cells PA levels are comparable to WT PR8, ruling out insufficient template availability or lower general translation efficiency as reasons for decreased PA-X activity. It is formally possible that NS1 has direct effect on PA-X activity in the nucleus or the cytoplasm, as both proteins are functioning in both of these compartments [7,25,27,68].

There are several limitations to our study. First, we intentionally utilized a MAVS-deficient cell line to allow for optimal replication and viral protein synthesis when NS1 is mutated. One of the important functions of host shutoff is to limit expression of antiviral genes. Our results demonstrate comparable host shutoff phenotypes in parental A549 cells and A549-ΔMAVS cells, however, as expected, we do not detect IFN or ISG transcripts in our nanopore sequencing reads. We also used a single laboratory adapted IAV strain PR8 and a single cell line. PR8 strain and A549 infection model are widely used in IAV research, and our results can be directly compared to other studies that use this model. However, it remains to be seen whether our findings hold true in other infection models.

## MATERIALS AND METHODS

### Cells

A549 (American Type Culture Collection (ATCC), Manassas, VA, USA) and A549-ΔMAVS cells [48] were maintained in Dulbecco’s modified Eagle’s medium (DMEM), supplemented with heat-inactivated 10 % fetal bovine serum (FBS) and 2 mM L-glutamine (all purchased from Thermo Fisher Scientific, Waltham, MA, USA) at 37 °C in 5 % CO₂ atmosphere.

### Viruses and infections

Wild-type A/PuertoRico/8/34(H1N1) (PR8-WT) and the mutant recombinant viruses PR8-PA(fs), PR8-NS1(N80) and PR8-PA(fs),NS1(N80) were generated as previously described [26]. Mutant recombinant viruses PR8-NS1(123A,124A) and PR8-NS1(187R) were generated using PCR site directed mutagenesis and the 8-plasmid reverse genetic system [69] as described previously [30], mutagenesis primer and plasmid sequences are available upon request. Virus stocks were produced in African green monkey kidney (Vero) cells. Influenza A virus strain A/California/7/09(H1N1) and influenza B virus strain B/Brisbane/60/08 were provided by the Public Health Agency of Canada (PHAC) National Microbiology Laboratory (Winnipeg, Canada) and propagated in Madin Darby Canine Kidney (MDCK) cells. Titers of all viral stocks were determined by plaque assays in MDCK cells using Avicel overlays as described in [70]. For each infection, cell monolayers were inoculated at multiplicity of infection (MOI) of 1 for 1 h at 37 °C. Then cells were washed briefly with Phosphate Buffered Saline (PBS, Thermo Fisher Scientific, Waltham, MA, USA) and cultured in infection medium (0.5 % Bovine Serum Albumin (BSA, Sigma-Aldrich, Missouri, USA) in DMEM) at 37 °C, 5 % CO_2_ atmosphere until the specified time post-infection. Vero and MDCK cells were obtained from ATCC (Manassas, VA, USA).

### Immunofluorescence staining

Cell fixation and immunofluorescence staining were performed according to the procedure described in [71]. Briefly, cells grown on 18-mm round coverslips were fixed with 4% paraformaldehyde in PBS for 15 min at ambient temperature and permeabilized with cold methanol for 10 min. After 1-h blocking with 5% bovine serum albumin (BSA, BioShop, Burlington, ON, Canada) in PBS, staining was performed overnight at +4⁰C with antibodies to the following targets: influenza A virus (IAV) polyclonal antibody (1:400, goat, Abcam, ab20841), NP (IAV) (1:1000, mouse, Santa Cruz, sc-101352), NP (IBV) (1:200, mouse, Santa Cruz Biotechnology, sc-57885), PABPC1 (1:1000, rabbit, Abcam, ab21060), PABPN1 (1:200, rabbit, Abcam, ab75855), SR proteins (1:100, mouse, Santa Cruz Biotechnology, sc-13509). Alexa Fluor (AF)-conjugated secondary antibodies used were: donkey anti-mouse IgG AF488 (Invitrogen, A21202), donkey anti-rabbit IgG AF555 (Invitrogen, A31572), donkey anti-goat IgG AF647 (Invitrogen, A32839). Where indicated, nuclei were stained with Hoechst 33342 dye (Invitrogen, H3570). Slides were mounted with ProLong Gold Antifade Mountant (Thermo Fisher) and imaged using Zeiss AxioImager Z2 fluorescence microscope or Leica TCS SP8 Confocal microscope. Green, red, blue, and far-red channel colors were changed for image presentation in the color-blind safe palette without altering signal levels.

### Single molecule fluorescent *in situ* hybridization (smFISH)

Cells grown on 18-mm round glass coverslips were briefly washed with PBS and fixed with 4 % paraformaldehyde in PBS for 10 min at room temperature. Permeabilization and hybridization steps were performed according to LGC Biosearch Technologies Stellaris RNA FISH protocol for adherent cells using human MALAT1 Stellaris FISH probe set with Quazar 570 dye (cat. Number SMF-2035-1), a custom Stellaris FISH probe set for IAV segment 7 genomic RNA (IAVM) with Fluorescein Dye (cat. number SMF-1025-5), or the 100 nM Alexa Fluor 555 labeled oligo-dT-40 probe (Thermo Fisher Scientific, Waltham, MA, USA). Nuclei were stained with Hoechst 33342 dye (Invitrogen, H3570). Glass coverslips were mounted with ProLong Gold Antifade Mountant (Thermo Fisher Scientific, Waltham, MA, USA) and imaged using Zeiss AxioImager Z2 fluorescence microscope or Leica TCS SP8 Confocal microscope. Green, red, blue, and far-red channel colors were changed for image presentation in the color-blind safe palette without altering signal levels.

### smFISH coupled to Immunofluorescence staining (ImmunoFISH)

Cells were processed for smFISH as described above before the coverslip mounting step, then washed with PBS for 5 min at room temperature. After 30 min blocking with 5% BSA in PBS, staining was performed overnight at + 4 °C with antibodies as described in the immunofluorescence staining section. Glass coverslips were mounted with ProLong Gold Antifade Mountant (Thermo Fisher Scientific, Waltham, MA, USA) and imaged using Zeiss AxioImager Z2 fluorescence microscope or Leica TCS SP8 Confocal microscope. Green, red, blue, and far-red channel colors were changed for image presentation in the color-blind safe palette without altering signal levels.

### Western Blotting

Whole-cell lysates were prepared by direct lysis of PBS-washed cell monolayers with 1× Laemmli sample buffer (50 mM Tris-HCl pH 6.8, 10% glycerol, 2% SDS, 100 mM DTT, 0.005% Bromophenol Blue). Lysates were immediately placed on ice, homogenized by passing through a 21-gauge needle, and stored at −20⁰C. Aliquots of lysates thawed on ice were incubated at 95⁰C for 3 min, cooled on ice, separated using denaturing PAGE, transferred onto PVDF membranes using Trans Blot Turbo Transfer System with RTA Transfer Packs (BioRad Laboratories, Hercules, CA, USA) according to manufacturer’s protocol and analyzed by immunoblotting using antibody-specific protocols. Antibodies to the following targets were used: actin (1:2000, HRP-conjugated, mouse, Santa Cruz Biotechnology, sc-47778), IFIT1 (1:1000, rabbit, Cell Signaling, #14769), influenza A virus (IAV) polyclonal antibody (1:2000, goat, Abcam, ab20841), ISG15 (1:1000, mouse, Santa Cruz, sc-166755), MAVS (1:1000, rabbit, Cell Signaling, #24930), NS1 (1:1000, mouse, clone 13D8, a gift from Kevin Coombs, [72]), PA (1:1000, rabbit, GeneTex, GTX125932), PABPN1 (1:1000, rabbit, Abcam, ab75855). For band visualization, HRP-conjugated anti-rabbit IgG (Goat, Cell Signaling, #7074) or anti-mouse IgG (Horse, Cell Signaling, #7076) were used with Clarity Western ECL Substrate on the ChemiDoc Touch Imaging Sysytem (Bio-Rad Laboratories). For analyses of protein band intensities, western blot signals were quantified using Bio-Rad Image Lab 5.2.1 software.

### RNA extraction and RT-qPCR

Total RNA was extracted using the RNeasy Plus (Qiagen, Hilden, Germany) kit protocol according to the manufacturer instructions. 250 ng of total RNA was used to synthesize cDNA using LunaScript®RT SuperMix (New England BioLabs Inc, Massachusetts, USA). Quantitative PCR amplification was performed using PerfeCta SYBR Green PCR master mix (QuantaBio, Beverly, MA, USA) and specific primers listed below on Cielo 3 QPCR unit (Azure Biosystems, California, USA). Primers used: MT CYB Left: cctaccctctcaacgacagc, MT CYB-Right: ctctgaccttttgccaggag, ACTB-Left: catccgcaaagacctgtacg, ACTB-Right: cctgcttgctgatccacatc; G6PD-Left: tgaggaccagatctaccgca, G6PD-Right: aaggtgaggataacgcaggc; POLR2A-Left: gaaacggtggacgtgcttat, POLR2A-Right: tgctgaaccaaagaacatgc; MALAT1-Left: gacggaggttgagatgaagc; MALAT1-Right: attcggggctctgtagtcct; PA-left: tctcagcggtccaaattcct; PA-right: tctgccagtacttgcttcca. Relative target levels were determined using ΔΔCt method using MT CYB as normalizer.

### Nuclear and cytoplasmic RNA fractionation

A549-ΔMAVS cells grown in 35-mm wells of 6-well cluster dishes were harvested at 24 hpi. Monolayers were placed on ice, briefly washed with ice-cold PBS, and incubated with 175 µL pre-chilled cytoplasmic lysis buffer [50 mM TrisCl pH 7.4, 1.5 mM MgCl₂, 140 mM NaCl, 0.5% IGEPAL (NP-40 substitute), 1 mM DTT, and 1U of RNaseOUT inhibitor]. Plates were incubated 5 min on ice. The cytoplasmic lysate was collected and resuspended with 350 µL of RLT+ buffer from RNeasy Plus kit (Qiagen, Hilden, Germany). Then, the nuclei were washed once with ice-cold cytoplasmic lysis buffer for 5 min on ice, buffer was removed, and nuclei were lysed in 350 µL of RLT+ buffer. Both lysates were mixed thoroughly by vortexing. Cytoplasmic and Nuclear RNA were extracted using the RNeasy Plus kit protocol according to the manufacturer’s instructions.

### MinION library preparation and sequencing

For each condition, poly(A) RNA isolation from Mock and IAV-infected A549-ΔMAVS cells was performed on a pool of six independent biological replicates. 20 µg of total RNA was diluted with nuclease-free water to a final volume of 200 µL and the poly(A) RNAs were isolated using the NEBNext Poly(A) mRNA Magnetic Isolation Module protocol (#E7490, New England BioLabs Inc, Massachusetts, USA). Nuclear and Cytoplasmic poly(A) RNA Libraries were prepared using the Oxford Nanopore Technology (ONT, Oxford, UK) Direct RNA Sequencing kit (SQK-RNA002) following the manufacturer’s protocol. In all libraries, 50 ng of poly(A) RNAs were used. Each final library was quantified using the Qbit 1X HS assay kit (Thermo Fisher Scientific, Massachusetts, USA). A total of 20 ng of prepared nuclear or cytoplasmic RNA library was loaded the same day onto a separate MinION R9.4 SpotON flow cell (FLO-MIN106) according to ONT specifications. The sequencing was run via MinKNOW software (v1.7.14) without live basecalling.

### MinION Bioinformatic processing

The FAST5 files were basecalled using Guppy (v3.2.4, ONT). The ONT long-read sequencing technology produces reads that are potentially full-length transcripts. Consequently, were not assembled; rather each read was treated as a complete transcript and used as such in the following analyses. The poly(A) tail lengths of the reads were estimated using Nanopolish-polyA (v10.2) (https://nanopolish.readthedocs.io/en/latest/quickstart_polya.html, [53]) on the reads previously aligned to the GRCh38.p13 human genome and PR8 reference genome using Minimap2 (v2.12) (https://github.com/lh3/minimap2; [73]) in splice mode. We used the mapping tool Isoquant (v3.3) (https://www.gencodegenes.org/human/, [74]) to assign a human gene to each human read using the GRCh38.p13 human genome and the human gene database gencode.v42. A version with and a version without mitochondrial sequences of both the genes and the genome references were used to delete reads for mitochondrial RNA. The Nanopolish-polyA and the Isoquant results were then combined to compute the average poly(A) length for each human gene. The Basic Local Alignment Search Tool BLASTN [75] was used to assign each viral read to one of 10 major viral transcripts, and these results were combined with the Nanopolish-polyA results to calculate an average poly(A) length for each viral transcript.

### Statistical analyses

Statistical analyses are described in figure legends. Analyses were performed using GraphPad Prism 9 software. For all datasets, N refers to the number of independent biological replicates performed on separate days.

## ACKNOWLEDGMENTS

We thank Richard Webby (St. Jude Children’s Research Hospital), Todd Hatchette (Dalhousie University), Kevin Coombs (University of Manitoba), and Alyson Kelvin (VIDO/University of Saskatchewan) for reagents. We also thank Dr. Gerard Gaspard and the Dalhousie CORES Cellular Microscopy and Digital Imaging Facility for assistance with fluorescence microscopy imaging.

## SUPPORTING INFORMATION CAPTIONS

**S1 Fig. Direct RNA sequencing of nuclear and cytoplasmic poly(A) RNAs using nanopore technology.** (A) Total RNA concentrations in nuclear and cytoplasmic fractions obtained from cell infected with the indicated recombinant mutant viruses or mock infected. Mean values from 6 independent replicates are plotted. Error bars represent standard deviations. (B) Summary of nanopore reads analyses obtained from poly(A) RNAs isolated from the cells infected with the indicated recombinant mutant viruses or mock infected. In each sample, yeast ENO2 spike in control RNA with 30 nt long poly(A) tail was added for comparison. H.sap = Homo sapiens; S.cer = Saccharomyces cerevisiae. (C) Human poly(A) transcripts identified in samples from WT PR8 virus-infected cells and absent in mock-infected cell samples. Nuc. = detected only in nuclear RNA sample (red); Nuc. & Cyt. = detected in both nuclear and cytoplasmic samples (orange); Cyt. = detected only in cytoplasmic sample (green).

**S1 Table. Read and poly(A) analysis summary for individual host transcripts.**

**S2 Table. Read and poly(A) analysis summary for individual viral transcripts.**

